# Evaluating the stoichiometric trait distributions of cultured bacterial populations and uncultured microbial communities

**DOI:** 10.1101/550681

**Authors:** Michael Manzella, Roy Geiss, E.K. Hall

**Author notes:** All work presented in this paper was conducted at Colorado State University, Fort Collins, CO USA.

## Abstract

The ecological stoichiometry of microbial biomass has most often focused on the ratio of the biologically-important elements carbon (C), nitrogen (N), and phosphorus (P) and has primarily been examined at a resolution where the contribution of the individual is masked by the reported population or community average. However, reporting population or community averages makes it difficult to assess phenotypic plasticity and stochasticity and mask important information required to understand both the drivers and implications of microbial biomass stoichiometry in nature. One way to assess the diversity of individual microbial phenotypes is through the use of single-cell techniques such as energy dispersive spectroscopy (EDS). EDS reports cellular quotas for the majority of elements composing microbial biomass including C, N, and P. In this study, by measuring individual cells within a microbial community or population, we describe for the first time the stoichiometry of microbial biomass as a distribution instead of an average. Exploration of stoichiometric trait distributions (as presented here) has the potential to improve our understanding of how nutrients interact with individual microorganisms to structure the elemental content of bacterial biomass and better describe how bacterial community biomass affects the ecosystems within which these organisms exist.

**Summary:** To assess the potential for EDS to describe the stoichiometric variance within populations and communities we measured the stoichiometric trait distribution of cultured freshwater bacterial populations under different resource conditions and compared them to natural microbial communities sampled from three lakes. Mean biomass C:N:P values obtained by EDS matched closely to those obtained by bulk measures using traditional analytical techniques for each freshwater isolate. However, we observed pronounced differences in the stoichiometric trait distributions of freshwater bacterial isolates compared to the stoichiometric trait distributions of natural communities. The stoichiometric trait distribution of the environmental isolates changed with P availability, growth phase, and genotype, with P availability having the strongest effect. The distribution of biomass ratios within each isolate growth experiment were the most constrained during stages of rapid growth and commonly had unimodal distributions. In contrast to the population distributions, the distribution of N:P and C:P for a similar number of cells from each of the mixed lake communities had narrower stoichiometric distributions and more commonly exhibited multiple modes.

## Introduction

Microorganisms are essential contributors to virtually all ecosystem processes. Individual microorganisms are selected to respond to environmental changes to optimize survival and reproductive success. Responses to the environment manifest as a suite of measurable morphological, physiological, and behavioral characteristics referred to as traits^1–3^. The study of microbial traits (phenotypes of ecological import that may or may not be phylogenetically conserved) and their inclusion into ecosystem models provide a path to improve our understanding of microbial influences over system-level processes^4–7^. However, due to technological and methodological challenges, estimates of microbial traits often lack a consideration of the variation within microbial communities or populations. If individual microorganisms are what responds to environmental change and affect system-level processes, then assessing the variation of a trait for a microbial population or community has the potential to better link the individual organism to the system-level processes it influences.

The study of microorganisms as individuals, or microbial individuality, has begun to elucidate the differences between the microorganism and the population or community in which it resides. Advances in technology, such as the use of Raman spectroscopy and secondary-ion mass spectrometry^8^, have allowed for individual microorganisms from natural environments to be studied and have found dramatic variability in traits from microbes from the same phylogenetic group residing in the same environment^9–11^. The causes of this individuality have been proposed to include genetic variation, micro-environmental heterogeneity, regulatory differences, and the stochasticity of metabolic reactions that overlay a conserved genome^11–14^. This last possibility has garnered support from the repeated studies of clonal laboratory populations that present varied values for continuous microbial traits, referred to as ‘phenotypic noise’^11, 12, 15–17^. The advantage, or consequence, of this variability may lie in bet-hedging or facilitating the development of microbial interactions within the population to allow for a division of labor^11, 14, 18, 19^. Thus, the traits of the community or population may be community-aggregated traits, where the properties of the individual determine the properties of the community, or they may be emergent traits where the properties of the community or population are greater than the sum of their parts^20^. Through the study of individuality of microbial physiology we can begin to decipher which traits are community-aggregated or emergent, and the impact of the individual at the ecosystem level^7, 20^.

One important microbial trait, biomass stoichiometry, provides a robust link between microbial physiology and ecosystem processes^21^. Studies of microbial biomass routinely evaluate the elemental composition of environmental microorganisms and have focused on the elements carbon (C), nitrogen (N), and phosphorus (P) due to their important role in biogeochemical cycles and ecosystem processes^22–27^. Community biomass stoichiometry is a composite of the stoichiometry of each constituent population which is a composite of each individual microbe^27^. Individual microorganisms may vary in elemental content due to differences such as their growth rate^12, 28–29^, cell size^30^, activity^31^, or growth phase^32^. Each individual is then comprised of a suite of macromolecules that have constrained elemental ratios^22, 33^. Thus, changes in the macromolecular composition of individual cells, in response to environmental cues, alters their stoichiometries^26^ and this propagates from individuals to populations to drive changes in community biomass stoichiometry^34^. This view of microbial community biomass is intuitive^27^, but investigation of microbial biomass within this framework is limited by the empirical resolution of commonly used methods. These methods typically include the concentration of bulk populations or the collection of mixed communities on a filter prior to elemental analysis using a total phosphorus spectrophotometric assay for P^35^ and a dry combustion method for C and N. Due to the limitation of measuring microbial biomass stoichiometry by wet chemistry, it has been primarily reported as a composite measure of a whole population or community. Regardless, bulk measurements of microbial biomass have provided important insights into the variation and function of this important community trait in nature and in the lab.

For example, it is relatively well documented that biomass C:N of aquatic bacteria is relatively conserved while both N:P and C:P biomass are more variable^36^. Resource stoichiometry of the surrounding environment, in particular P availability (e.g., resource C:P, hereafter C:P_R_), is a principal driver of biomass N:P and C:P^37–40^. Microbial responses to changing environmental conditions such as C:P_R_ results in substantial variability, or plasticity, of traits involving P^21^. This variability can be explained by the ability of microbial cells to alter the composition of macromolecules in their biomass to accommodate changing environmental and growth conditions^22, 32, 41–46^. For example, the growth rate hypothesis posits that P increases in rapidly-growing cells as increased growth rate is driven by an increase in nucleic acid-rich ribosomes which can be a principal pool of P in a cell^22^. The variability is exacerbated by the ability of microbial cells to store C-rich and P-rich compounds^47, 48^. Recent evidence also suggests that phylogeny likely plays a role in biomass stoichiometry^29, 38^, although biomass stoichiometry is rarely homeostatic even within a single isolate^38, 39–49^. Unlike biomass stoichiometry itself, there is no definitive evidence that stoichiometric plasticity has a phylogenetic signal^38^. When stoichiometry is examined in the laboratory, it appears as though the plasticity of biomass N:P and C:P is greater than that observed within natural communities^38^. This indicates that this plasticity may be an artifact of the laboratory conditions, and may not reflect realized *in situ* phenotypes. Whereas the relationship between microbial biomass and important environmental variables has been advanced through studies of bulk biomass^38, 50, 51^, due to the empirical constraints of measuring microbial biomass C:N:P we know little of how individuals contribute to the population and community mean that define these relationships.

One approach to increase our understanding of microbial biomass stoichiometry in both cultured populations and uncultured microbial communities is to employ single-cell stoichiometric analyses. Energy dispersive spectroscopy (EDS), also referred to as x-ray microanalysis (XRMA), is one such technique that has the potential to enable single-cell analyses. Early studies demonstrated the ability of EDS to analyze biomass stoichiometry in individual bacterial and algal cells^52, 53^. However, these studies commonly reported averaged trait values and did not present the distribution of these stoichiometric traits. Since these early studies, the equipment and techniques used for EDS have improved and enabled single-cell stoichiometries to be detected and reported with greater sensitivity and in higher resolution^54–56^. Here, we aim to advance the description of the stoichiometric trait distribution for cultured and uncultured bacteria by integrating single cell EDS-generated stoichiometric data into probability density functions to describe the frequency distribution of stoichiometric traits of individual bacterial cells within a population or community.

Probability density functions have been proposed as a possible best-practice to enable quantitative descriptions of trait diversity^1^. These probability density functions are analogous to Hutchinson’s niche concept^57^ where the niche space or in this case trait space can be represented by a probabilistic hypervolume^1^. Trait distributions can be expanded into several components which include richness, evenness, divergence; and, if we examine trait distribution between different systems, we can also estimate distinctiveness^1, 3^. When a trait distribution is represented as a probability density function, the richness is the breadth of reported phenotypes, evenness (as the name suggests) is how evenly distributed the phenotype is across the phenotypic space, and divergence is a measure of the distribution of trait abundance toward the extremes of the occupied phenotypic space^,13^. When trait distributions are compared among populations or communities, distinctiveness is analogous to β-diversity – a measure of the difference between the trait distribution of each group^1^. These trait distribution metrics provide a way to qualitatively and quantitatively describe how elements are distributed among cells, within populations, and among cells within communities cultured under different conditions or sampled from different ecosystems.

In this study we demonstrate the potential of EDS to describe the stoichiometric trait distribution of bacterial cultures grown in the lab under different resource conditions and natural microbial communities sampled directly from the environment. We first generated calibration factors from three biologically-relevant compounds to relate characteristic counts for C, N, and P to elemental molar equivalents. We validated these calibration factors by comparing the average stoichiometric ratios of three microbial populations estimated by EDS and traditional bulk measures. To determine how stoichiometric trait distributions were affected by genotype, C:PR, and by growth phase we analyzed three freshwater bacterial isolates at multiple points during a batch growth experiment at two C:PR levels. Finally, we analyzed the stoichiometric trait distributions of three natural microbial communities sampled from three Colorado lakes. These studies allowed us to compare the stoichiometric distributions from isolate populations (where genotype, growth phase, and resource conditions were known) to those of lake communities (where genotype, growth phase, and resource conditions were unknown).

## Results

### Comparison of EDS and bulk measurements of biomass stoichiometry

We first assessed the similarity of EDS measurements we generated to the more common community bulk measures of biomass stoichiometry. We compared the biomass C:N:P generated from averaged EDS-generated data (n > 90 cells, ea.) from three freshwater bacterial isolates grown on 100:1 C:P_R_ media to those obtained from the same samples after collection on pre-combusted GF/F filters (n = 3) and subsequent bulk determination of biomass stoichiometry (Figure 1 and Table 1). We found that the biomass C:N and C:P were not significantly different when examined by traditional bulk measures and EDS, with the exception of *Pseudomonas* UND-L18 (p = 0.004 and 0.023, respectively) which was enriched in C in the EDS-estimated elemental ratios when compared to bulk measures. Biomass N:P was not significantly different between techniques for any isolate (Figure 1 and Table 1).

**Figure 1:**
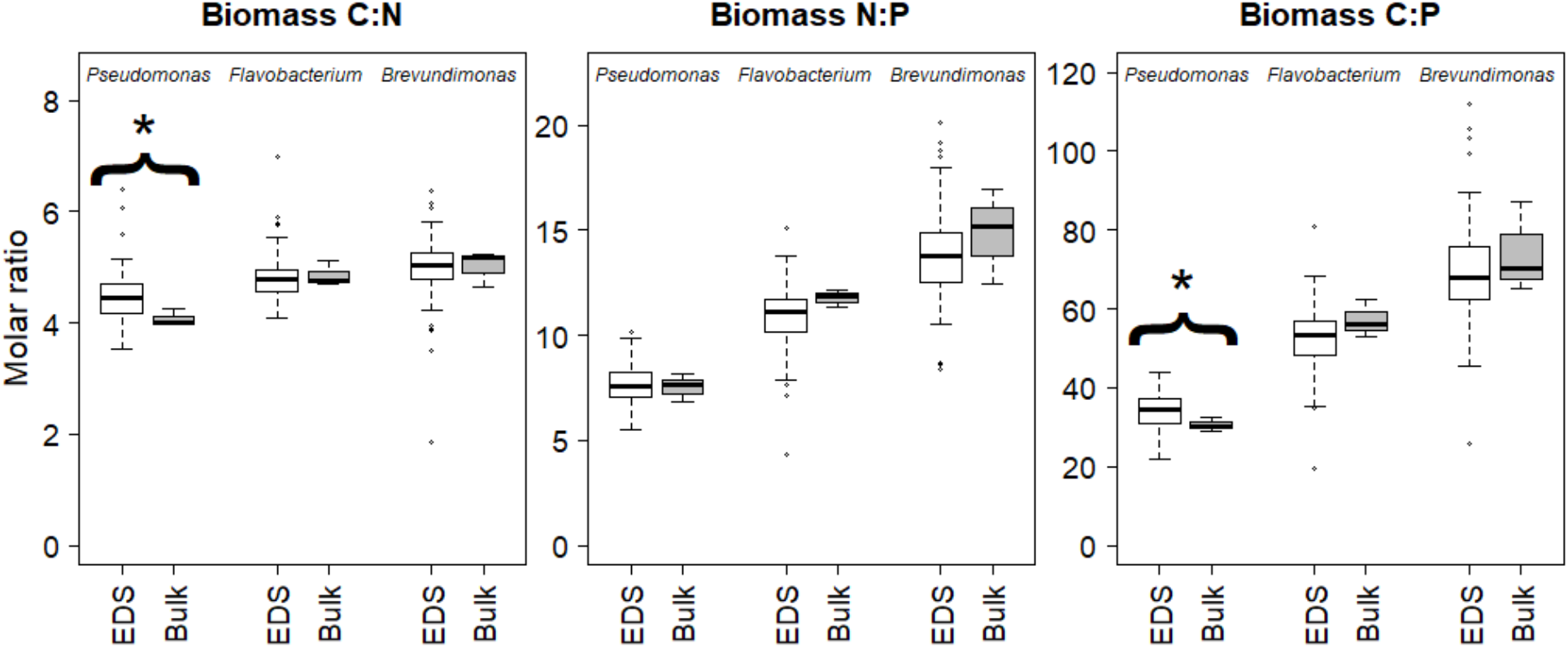
Stoichiometric trait values generated from EDS spectra (open boxes, n > 90 cells) compared to those obtained by bulk measures of stoichiometry (shaded boxes, n =3 filters). Asterisks indicate a significant difference (p < 0.05) between values obtained by EDS and the bulk measures.

**Table 1:** Biomass ratios generated by EDS and bulk measures of stoichiometry. Asterisks indicate that the biomass stoichiometry values by EDS were significantly different from those obtained by bulk measures (p ≤ 0.05).

| Isolate | Biomass C:N |  | Biomass N:P |  | Biomass C:P |  |
| --- | --- | --- | --- | --- | --- | --- |
|  | EDS | Bulk | EDS | Bulk | EDS | Bulk |
| *Pseudomonas* | 4.49 ± 0.46 | 4.07 ± 0.15* | 6.64 ± 0.93 | 7.53 ± 0.68 | 34.18 ± 4.54 | 30.61 ± 1.85* |
| *Flavobacterium* | 4.83 ± 0.40 | 4.86 ± 0.23 | 10.93 ± 1.47 | 11.77 ± 0.43 | 52.76 ± 8.09 | 57.76 ± 4.64 |
| *Brevundimonas* | 4.99 ± 0.48 | 5.01 ± 0.31 | 13.89 ± 2.00 | 14.84 ± 2.26 | 69.21 ± 12.04 | 74.27 ± 11.68 |

\*Indicates EDS and bulk measures were significantly different from one another ( $p \leq 0.05$ )

### Stoichiometric plasticity across a batch growth experiment

To explore the effect of growth phase on the stoichiometric trait distribution of bacterial biomass at an estimated non-limiting P resource ratio^36, 39^, we grew three freshwater bacterial isolates on 100:1 C:P_R_ medium. Soluble P of the liquid media began at or near the intended 239 μM P for each culture. In each case, soluble P rapidly decreased as optical density (OD) increased (Supplemental Figure 1). OD began to plateau in *Pseudomonas* UND-L18 at ~8 h as soluble P reached ~2-4 μM P; *Flavobacterium* N288 had reached stationary phase prior to P depletion at 15 h; and the OD of *Brevundimonas* N1445 reached stationary phase at a relatively enriched level of soluble P (~ 70 μM). Throughout the 100:1 C:P_R_ growth experiment, the biomass C:N, N:P, and C:P changed little across growth phase for each isolate (Figure 2 and Table 2). For all isolates, biomass C:N was the constrained whereas N:P and C:P showed minor but significant (p ≤ 0.05) decreases as cells entered exponential growth (Figure 2 and Table 2). Within each growth phase, biomass N:P and C:P for each isolate was significantly different from each other (p ≤ 0.05), with the exception of *Pseudomonas* UND-L18 and *Flavobacterium* N288 during lag phase. For all growth phases, *Pseudomonas* UND-L18 had the lowest N:P and C:P followed by *Flavobacterium* N288 and then *Brevundimonas* N1445 having the highest average N:P and C:P (Supplemental Table 1). This trend continued until the stationary-phase sampling where the N:P and C:P ratios of *Flavobacterium* N288 decreased to below those of *Pseudomonas* UND-L18.

**Figure 2:**
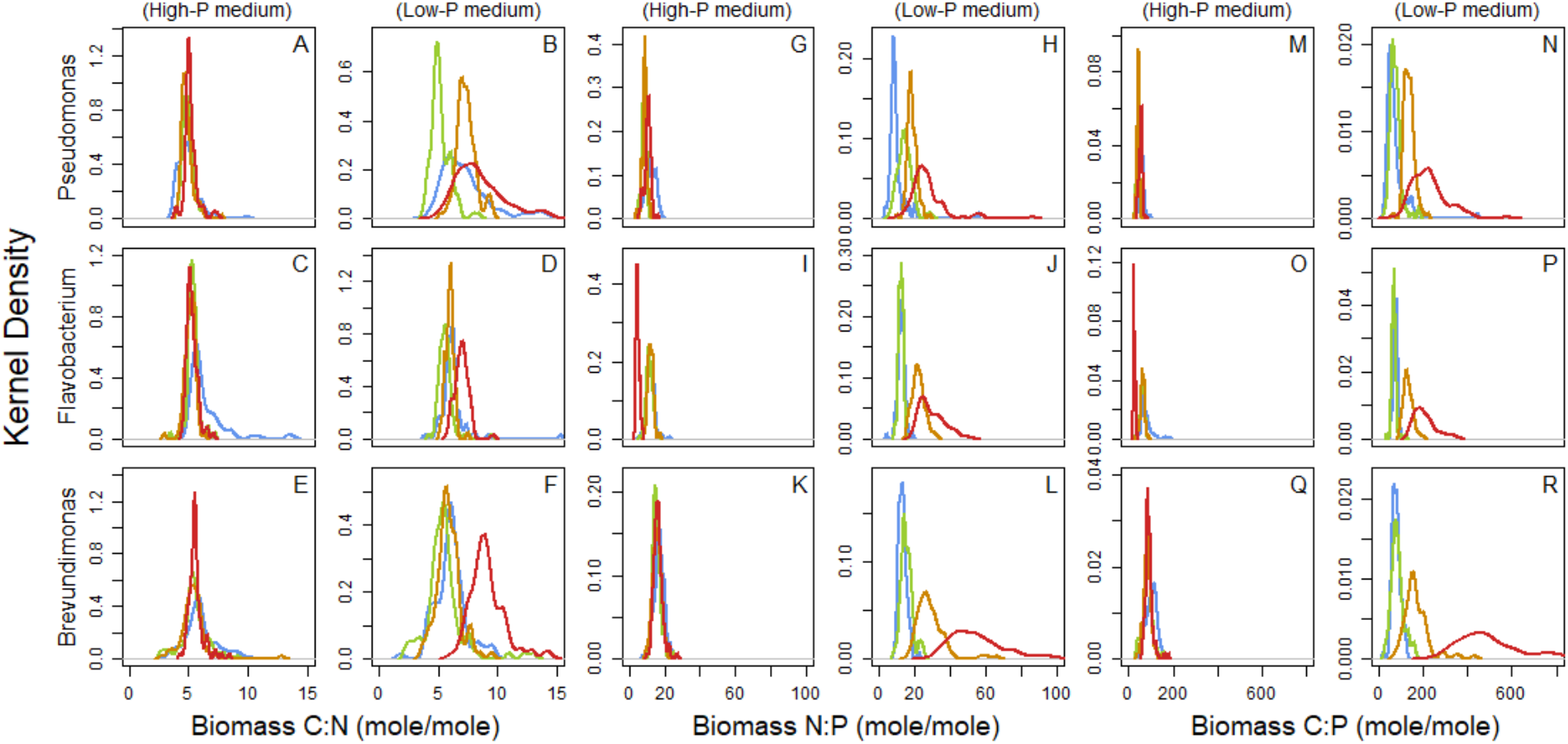
Stoichiometric trait distributions from three freshwater isolates grown on 100:1 and 1000:1 C:P_R_ media. Biomass C:N (panels A-F), biomass N:P (panels G-L), and biomass C:P (panels M-R) are normalized to show the plasticity of these traits across all media conditions and isolates. Growth phase is represented by colors with **blue** indicating lag, **green** indicating early-log, **orange** indicating mid-log, and **red** indicating stationary phase.

**Table 2:** Average population-level biomass trait values for three environmental isolates as measured at each phase of a batch growth experiment. The superscripts L, E, M, and S in dicate a significant difference (p ≤ 0.05) in the trait value when analyzed during lag (L), early-log (E), mid-log (M), and stationary phase (S), for the same resource C:P and isolate. Tables depicting statistical differences by isolate and C:P_R_ can be found in Supplemental Tables 2 and 3, respectively.

| *Pseudomonas* |  |  |  |  |  |
| --- | --- | --- | --- | --- | --- |
| Trait | Resource C:P | Lag | Early log | Mid log | Stationary |
| Biomass C:N | 100:1 | <sup>S</sup> 4.94 ± 0.90 | <sup>S</sup> 4.97 ± 0.45 | <sup>S</sup> 4.89 ± 0.53 | <sup>L, E, M</sup> 5.21 ± 0.55 |
|  | 1000:1 | <sup>E, S</sup> 7.44 ± 2.57 | <sup>L, M, S</sup> 5.20 ± 0.80 | <sup>E, S</sup> 7.35 ± 0.80 | <sup>L, E, M</sup> 8.47 ± 2.05 |
| Biomass N:P | 100:1 | <sup>E, M, S</sup> 12.39 ± 2.77 | <sup>L, M, S</sup> 8.61 ± 1.30 | <sup>L, E, S</sup> 9.03 ± 1.41 | <sup>L, E, M</sup> 10.71 ± 1.80 |
|  | 1000:1 | <sup>E, M, S</sup> 10.03 ± 5.80 | <sup>L, M, S</sup> 14.80 ± 4.23 | <sup>L, E, S</sup> 18.83 ± 2.69 | <sup>L, E, M</sup> 27.65 ± 10.80 |
| Biomass C:P | 100:1 | <sup>E, M, S</sup> 59.55 ± 9.82 | <sup>L, S</sup> 42.61 ± 6.06 | <sup>L, S</sup> 43.71 ± 5.47 | <sup>L, E, M</sup> 55.29 ± 7.71 |
|  | 1000:1 | <sup>M, S</sup> 73.85 ± 49.12 | <sup>M, S</sup> 76.82 ± 25.68 | <sup>L, E, S</sup> 138.33 ± 23.66 | <sup>L, E, M</sup> 228.84 ± 82.26 |

| *Flavobacterium* |  |  |  |  |  |
| --- | --- | --- | --- | --- | --- |
| Trait | Resource C:P | Lag | Early-log | Mid-log | Stationary |
| Biomass C:N | 100:1 | <sup>E, M, S</sup> 6.41 ± 1.54 | <sup>E, M</sup> 5.36 ± 0.48 | <sup>L, E, M</sup> 5.15 ± 0.45 | <sup>L, M</sup> 5.31 ± 0.45 |
|  | 1000:1 | <sup>E, S</sup> 6.22 ± 1.20 | <sup>L, M, S</sup> 5.61 ± 0.65 | <sup>E, S</sup> 5.98 ± 0.35 | <sup>L, E, M</sup> 6.96 ± 0.59 |
| Biomass N:P | 100:1 | <sup>E, S</sup> 12.43 ± 2.36 | <sup>L, M, S</sup> 11.62 ± 1.82 | <sup>E, S</sup> 12.41 ± 1.50 | <sup>L, E, M</sup> 5.12 ± 0.95 |
|  | 1000:1 | <sup>M, S</sup> 12.72 ± 2.00 | <sup>M, S</sup> 12.51 ± 1.27 | <sup>L, E, S</sup> 22.38 ± 3.48 | <sup>L, E, M</sup> 29.70 ± 6.90 |
| Biomass C:P | 100:1 | <sup>E, M, S</sup> 79.22 ± 22.96 | <sup>L, S</sup> 62.23 ± 10.15 | <sup>L, S</sup> 64.06 ± 9.33 | <sup>L, E, M</sup> 27.00 ± 4.24 |
|  | 1000:1 | <sup>E, M, S</sup> 77.63 ± 8.89 | <sup>L, M, S</sup> 70.18 ± 10.70 | <sup>L, E, S</sup> 133.85 ± 21.09 | <sup>L, E, M</sup> 206.03 ± 48.23 |

**Table 2:** Average population-level biomass trait values for three environmental isolates as measured at each phase of a batch growth experiment.
| *Brevundimonas* |  |  |  |  |  |
| --- | --- | --- | --- | --- | --- |
| Trait | Resource C:P | Lag | Early-log | Mid-log | Stationary |
| Biomass C:N | 100:1 | <sup>E, M, S</sup> 6.02 ± 1.34 | <sup>L</sup> 5.47 ± 1.10 | <sup>L</sup> 5.39 ± 1.17 | <sup>L</sup> 5.63 ± 0.63 |
|  | 1000:1 | <sup>E, S</sup> 6.04 ± 1.29 | <sup>L, S</sup> 5.50 ± 1.66 | <sup>S</sup> 5.88 ± 1.00 | <sup>L, E, M</sup> 9.14 ± 1.50 |
| Biomass N:P | 100:1 | <sup>E, M</sup> 17.20 ± 3.01 | <sup>L, S</sup> 15.75 ± 2.32 | <sup>L</sup> 15.97 ± 2.62 | <sup>E</sup> 16.52 ± 2.51 |
|  | 1000:1 | <sup>E, M, S</sup> 13.50 ± 2.25 | <sup>E, M, S</sup> 15.78 ± 3.21 | <sup>L, E, S</sup> 29.06 ± 8.07 | <sup>L, E, M</sup> 56.35 ± 17.96 |
| Biomass C:P | 100:1 | <sup>E, M, S</sup> 102.47 ± 24.37 | <sup>L, S</sup> 86.41 ± 22.23 | <sup>L, S</sup> 85.04 ± 17.01 | <sup>L, E, M</sup> 92.83 ± 17.46 |
|  | 1000:1 | <sup>M, S</sup> 80.80 ± 18.68 | <sup>M, S</sup> 86.37 ± 28.36 | <sup>L, E, S</sup> 171.06 ± 57.15 | <sup>L, E, M</sup> 510.37 ± 165.64 |

The differences in biomass stoichiometry among isolates and growth phases noted above are based on averaged trait values for *ca.* 90 cells, yet the benefit of EDS-based stoichiometric measurements is their integration into trait distributions and the trait framework. By examining the stoichiometric trait distributions for these isolates, we note that the decrease in average biomass N:P and C:P during log phase is accompanied by a general decrease in richness and divergence for each trait (Table 3). The majority of the trait distributions for biomass C:N, N:P, and C:P from the isolate experiments were unimodal where the average trait value corresponded to the distribution maximum (Figure 2 and Table 2). In addition, we note that the richness of *Flavobacterium* N288 was the lowest across each stoichiometric ratio, followed by *Pseudomonas* UND-L18, and finally *Brevundimonas* N1445 that was the richest and subsequently exhibited the greatest stoichiometric plasticity (Figure 2 and Tables 2–3). Evenness showed no clear trends among isolates, growth phase, or between media resource ratios (Table 3).

**Table 3:**
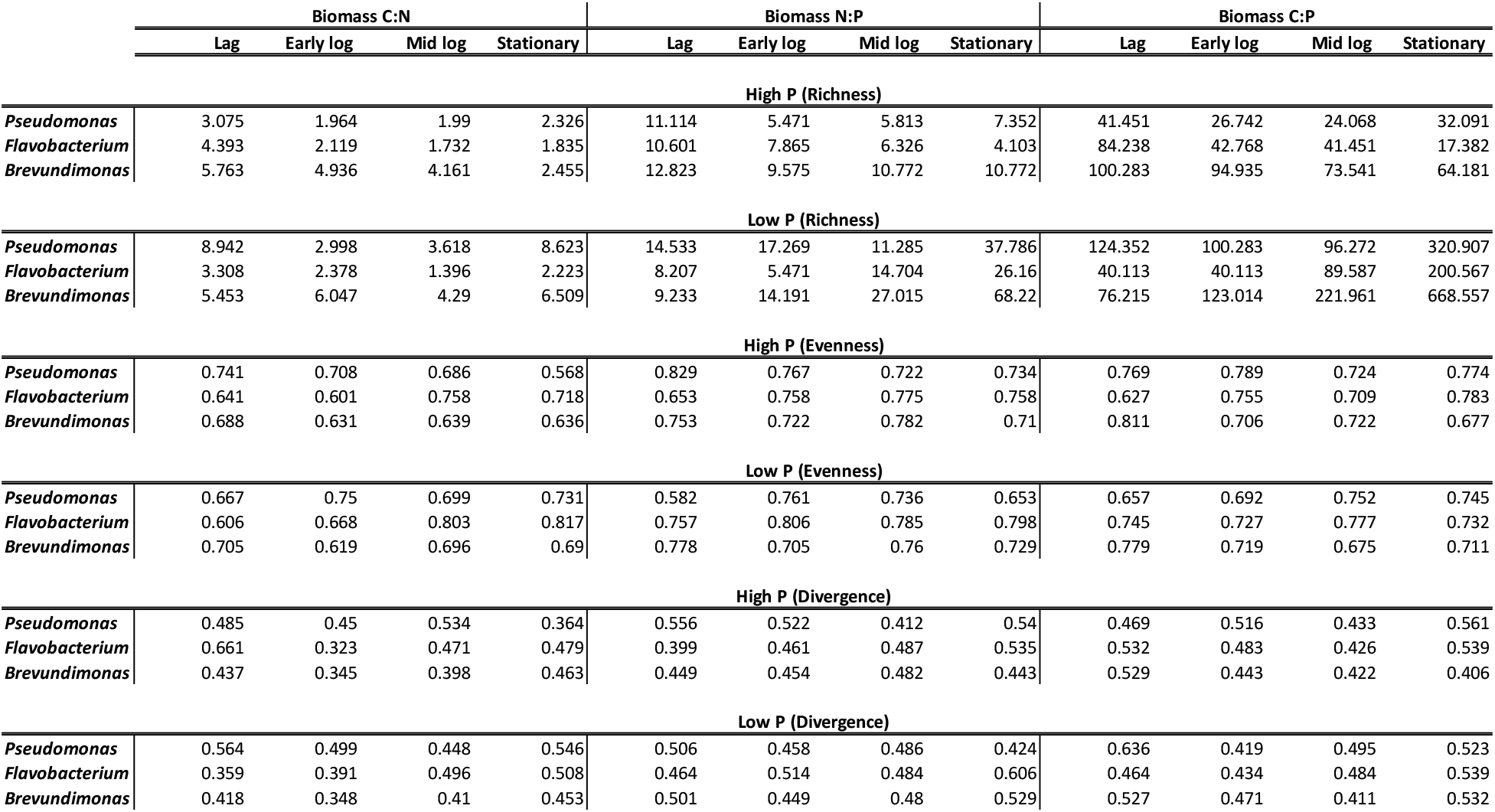
Characteristics of trait diversity for the isolate experiment sshowing trait richness, evenness, and divergence across three freshwater isolates, four growth phases, and two C:P_R_ ratios.

We repeated each of the above growth experiments using a low-P growth medium (C:P_R_ = 1000) to assess the dynamics of stoichiometric trait distributions in a more P-limited environment^36, 39^ Resource P in the second set of experiments began near the intended 23.9 μM P. As in the 100:1 C:P_R_ medium, P rapidly decreased as the OD of each culture increased (Supplemental Figure 1). *Pseudomonas* UND-L18 removed >98 % of the soluble P (to *ca.* 0.3 μM P) within 5 hours whereas *Flavobacterium* N288 and *Brevundimonas* N1445 took 10 h and 15 h to deplete the soluble P to the same level, respectively. In the low-P growth experiments, OD continued to increase after P depletion until the final time point (55 hours); at which time we sacrificed the cultures. Between growth phases, we saw significant changes in the average stoichiometry (p ≤ 0.05), especially in biomass C:P and N:P in early- and mid-log phase (Figure 2 and Table 2). Biomass N:P and C:P for each isolate increased significantly (p ≤ 0.05) throughout the experiment (Table 2), with *Brevundimonas* N1445 again exhibiting the greatest stoichiometric plasticity (Figure 2 and Table 2). In addition, biomass C:N was significantly higher during stationary phase than any other growth phase for each examined culture (Table 2). In the low-P media, we detected significant differences in biomass C:N, N:P, and C:P across the three isolates at each sampling point with the exception of biomass C:P during lag phase where the three isolates were not significantly different (Table 2 and Supplemental Table 1). In addition, there were significant differences between the average trait values within a single genotype when grown on the 100:1 and 1000:1 C:P_R_ growth media (Supplemental Table 2) during some but not all phases of growth.

As in the 100:1 C:P_R_ cultures, the stoichiometric trait distributions from the 1000:1 C:P_R_ cultures were primarily unimodal and corresponded to the community average (Figure 2 and Table 2). Phenotypic richness and divergence (Figure 2 and Table 3) were generally lower during stages of rapid growth, in this case lag and early-log phases. As growth proceeded with media P near the detection limit, the richness of biomass N:P and C:P increased for all three isolates (Table 3). *Brevundimonas* N1445 again exhibited the greatest phenotypic richness for each trait, with the exception of biomass C:N where *Pseudomonas* UND-L18 had higher richness during lag- and stationary-phase growth (Figure 2 and Table 3). These 1000:1 C:P_R_ cultures defined the stoichiometric ranges used for the trait probability density classifications as we noted their stoichiometric richness was the greatest across the entirety of the study, including the analysis of the natural communities described below. We set the trait range for biomass C:N to 0 – 20, biomass N:P to 0 – 100, and biomass C:P to 0 – 1000 and used these ranges to assess the characteristic of each stoichiometric trait distribution that we measured.

### Analysis of uncultured microbial cells

To assess how the stoichiometric trait distribution of genotypicaly-defined cells (in a known physiological state and defined media) compared to diverse communities (in an unknown physiological state and undefined resource conditions), we analyzed natural microbial communities from three small lakes in Colorado (Loch Vale, Warren Lake, and Sheldon Pond). Loch Vale is an oligotrophic (mean ice-free Chl *a* = 2.2 +/− 1.9 μg L^−1^) alpine lake whereas Warren Lake and Sheldon Pond are both urban, mesotrophic lakes (mean ice-free Chl *a* =7.9 +/− 0.9 μg L^−1^ and 12.3 +/−0.8 μg L^−1^, respectively). Average microbial community biomass C:N was similar among each lake ecosystem (Figure 3 and Table 4), although Warren Lake had significantly higher biomass C:N than both Loch Vale and Sheldon Pond (p ≤ 0. 005). In addition, richness was low for biomass C:N, with each stoichiometric trait distribution occupying under 15 % of the defined phenotypic landscape (0-20 biomass C:N) for all lake communities (Figure 3 and Table 4). Evenness for biomass C:N was generally higher in the lake communities than in the isolate experiments while divergence was lower (Tables 3 and 4). In addition, the stoichiometric trait distribution for C:N was unimodal for all lake communities (Figure 3).

**Figure 3:**
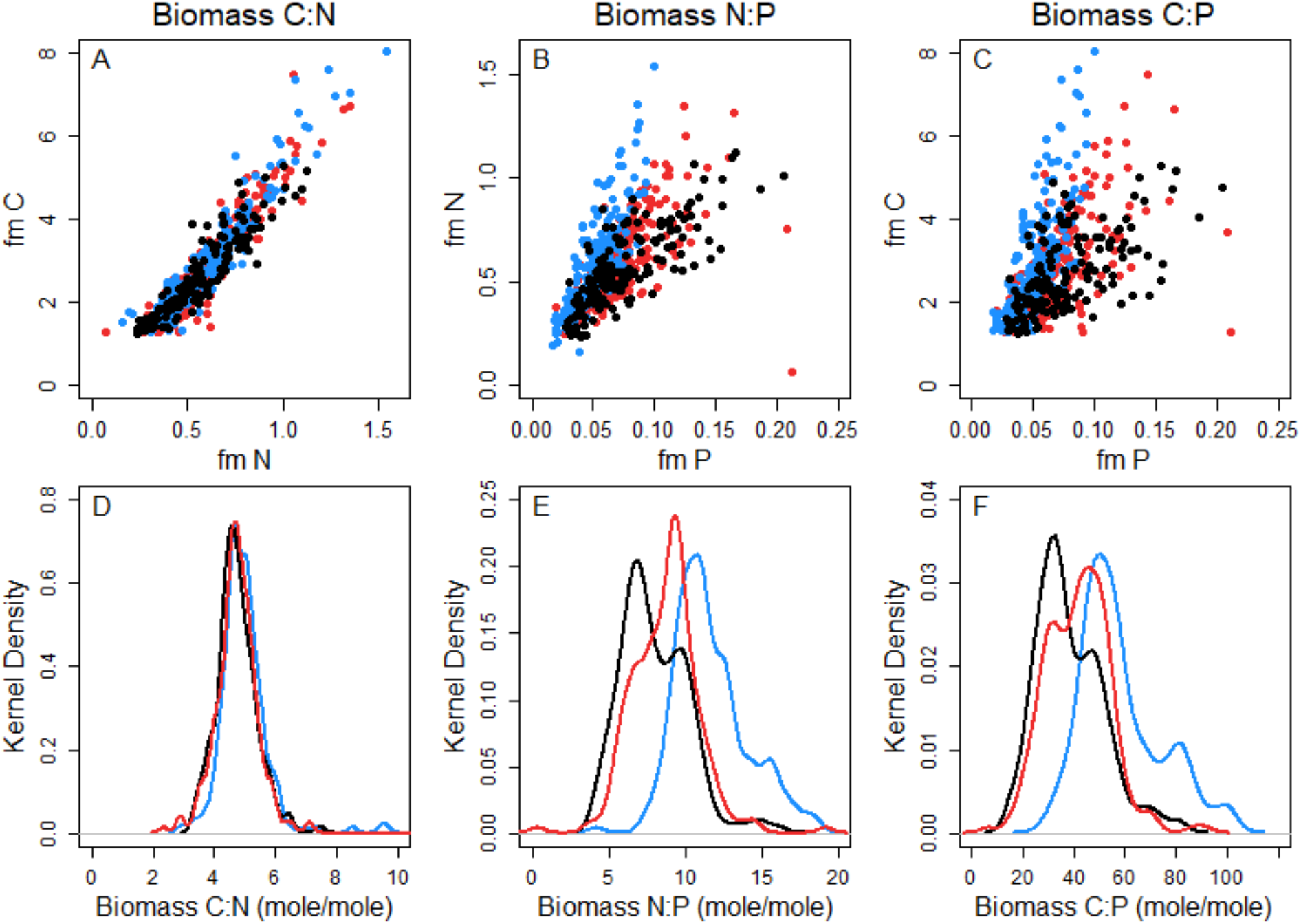
Stoichiometric trait distributions from three lake communities. Distributions are color coded by lake community with **red** representing Loch Vale, **blue** representing Warren Lake, and **black** representing Sheldon Pond.

**Table 4:**
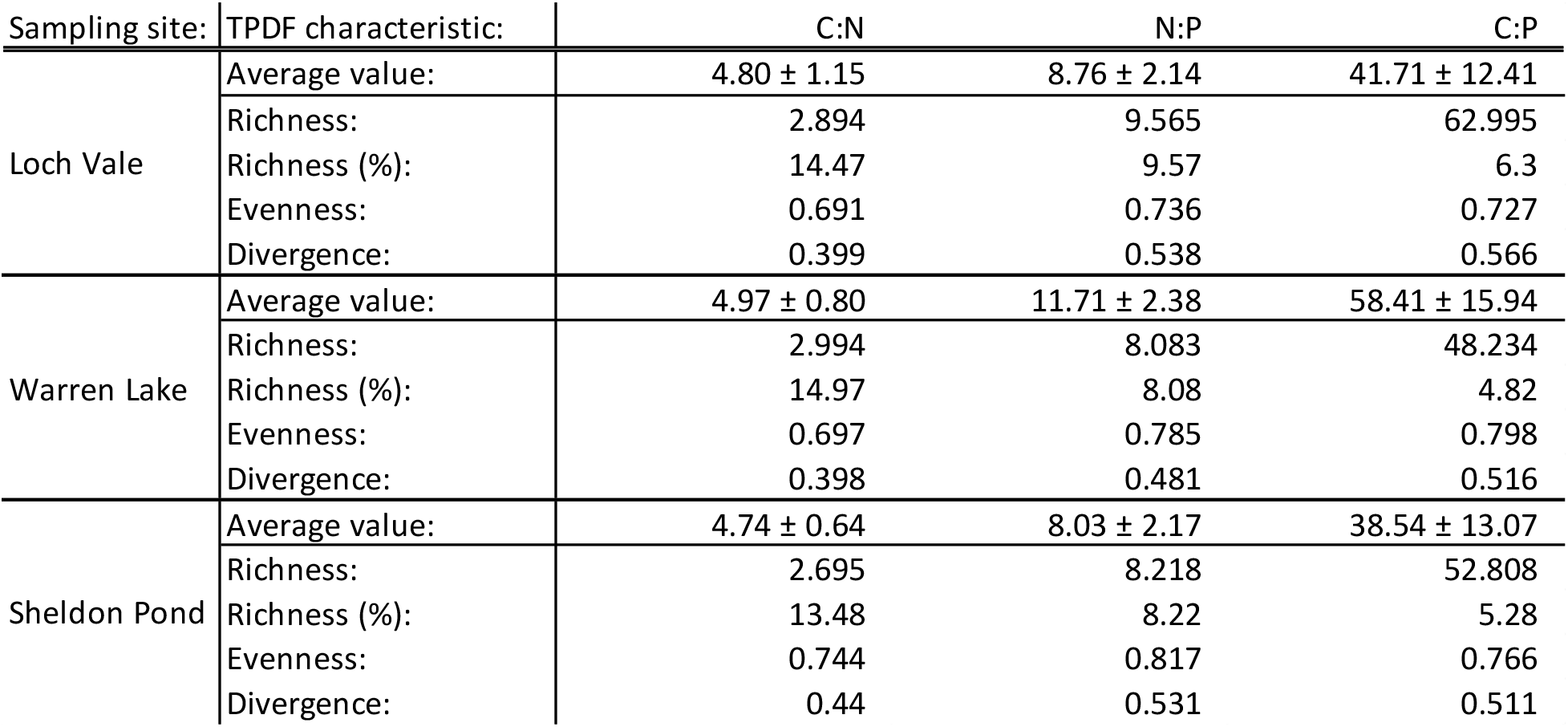
Characteristics of the stoichiometric trait distribution for each stoichiometric trait across three lakes as calculated by the TPD package in the R software platform. We analyzed each strain, time point, and lake community simultaneously and selected the default alpha value of 0.95 for each trait. We set the trait range for biomass C:N to 0 – 20, biomass N:P to 0 – 100, and biomass C:P to 0 – 1000. We set the number of divisions within those trait ranges to provide one division per 0.1 change in molar ratio.

We predicted that biomass P and its associated traits (N:P and C:P) would vary by lake and have increased richness and divergence compared to the isolate experiments. The microbial communities from Loch Vale, Warren Lake, and Sheldon Pond all showed significant differences in biomass N:P and C:P from one another (p ≤ 0.005) (Table 4). Unlike the isolate experiments where the majority of N:P and C:P distributions were unimodal, the stoichiometric trait distributions for N:P and C:P from these mixed communities were multimodal with local maxima that were distinct from the community mean biomass C:P and N:P (Figure 3 and Table 4). For instance, the Warren Lake microbial community had a C:P local maximum of ~50 with additional (less abundant but clearly defined) local maxima at ~82 and ~99. The same was true in the Loch Vale community which had a local maximum for biomass C:P of ~46 with an additional local maximum at C:P of ~32. Interestingly, the Sheldon Pond community had local maxima at or near those of Loch Vale (~33 and ~47), although the primary and secondary modes were inverted compared to Loch Vale (Figure 3). Whereas the biomass C:N richness was similar between lake communities and isolate populations during rapid growth, biomass N:P and C:P within the lake communities were, on average, lower in richness (i.e. more constrained) compared to the environmental isolates grown on the low-P media (Tables 3 and 4). Thus, the stoichiometric trait distributions of natural communities differed from those of the natural populations by being more constrained (i.e. lower richness) and were multi-modal compared to the unimodal distributions that were characteristic of the isolate growth experiments.

## Discussion

In this study we report the distribution that underlies an average community biomass stoichiometry of bacterial populations by using batch growth experiments to evaluate the relative influence of genotype, growth phase, and C:P_R_ on the stoichiometric trait distribution. We use the results of the batch culture experiments to compare the stoichiometric trait distributions generated from populations at known physiological states and cultivated under defined resource conditions to those generated from three natural lake communities where the physiological state and local resource conditions were unknown.

### Isolate experiments

In the high-P media (C:P_R_ = 100) we saw subtle but clear differences in the stoichiometric trait distribution among genotype and growth phase that are consistent with our existing understanding of drivers of biomass stoichiometry. For instance, the decreases in richness and divergence that occurred during early- and mid-log phase in the 100:1 C:P_R_ treatment for each isolate and each stoichiometric trait (Figure 2 and Table 3) are consistent with the previously reported idea that biomass composition becomes more constrained during rapid growth^28, 58^. In addition, the fastest growing cells (Supplemental Figure 2) exhibited the lowest values for biomass N:P and C:P (Table 2) which is consistent with the growth rate hypothesis^8^. In these monoculture experiments, the distribution of biomass values for each biomass trait was primarily unimodal, indicating that the biomass ratios existed around an optimal value for a given nutrient condition. Whereas we did detect significant changes in the average biomass stoichiometry and subtler differences in the stoichiometric trait distribution in the high-P media as the cultures expended the available P, the scale of these changes were smaller than those seen in the low-P cultures (Figure 2). This, coupled with the rapid plateau of OD values, may indicate that even at the single-cell level, cultures grown in an excess of available nutrients are unable or unprepared to shift their biomass ratios upon nutrient depletion. The 100:1 C:P_R_ growth medium also represents a C limited resource environment^39^, which may impose limits on bacterial growth not seen in the 1000:1 C:P_R_ growth medium. C-limitation has been shown to more abruptly halt bacterial replication compared to P depletion due to the role of C in catabolic metabolism whereas P is more closely linked to anabolic metabolism and required in smaller amounts^59, 60^.

P-limitation has been shown to result in increased stoichiometric plasticity of microbial biomass across a range of isolates^28, 39^. Under P-limiting conditions, we predicted and observed changes in the stoichiometric trait distribution of N:P and C:P, but we also found shifts in the trait distribution for biomass C:N when compared to C:N trait distribution for the low-P media (Figure 2 and Table 2). This is likely due to the disproportionate accumulation of C relative to N under P-limitation, as has been previously reported^38, 39^. In effect, cells under P-limitation uptake excess C and incorporate it into storage molecules^39, 60^. N storage is less common^21, 38^ and thus this is consistent with the increase in biomass C:N that we observed in the low-P media experiments. An increase in richness for both N:P and C:P biomass in the low-P experiments (Figure 2 and Table 3) indicated that the individuals within the population did not respond evenly or consistently to changing C:P_R_ which may be linked to bet-hedging or division of labor^14^. Yet, even as richness increased across growth phase (Figure 2 and Table 3), there was a pronounced difference in the average stoichiometry (Table 2) and stoichiometric trait distribution (Figure 2) for each isolate. This suggests that genotype has an important influence on the stoichiometric trait distributions. Additionally, the different genotypes appeared to respond differently by growth phase on the 100:1 and 1000:1 C:P_R_ media (Supplemental Table 2). This suggests differences in stoichiometric sensitivity when cells are growing near their maximum growth rate with abundant resources, as compared to growing more slowly under P-limiting conditions^28^. Whereas we only evaluated three isolates in this study, these results are consistent with previous studies that have demonstrated that genotype affects biomass stoichiometry between isolates grown on the same media^29, 38^.

The analysis of three environmental isolates grown under different resource conditions and sampled during different growth phases demonstrated the potential for significant intrapopulation phenotypic plasticity. However, in the vast majority of treatments (~80 %), these isolate experiments produced nearly unimodal stoichiometric trait distribution for biomass C:N, N:P, and C:P, even as richness increased as culatures entered stationary phase (Figure 2 and Table 3). In the remaining 15 cases we detected secondary modes, which we propose is the result of spatial and temporal heterogeneity present within the batch culture vessels where culture conditions were constantly changing. We predict that cultures grown in chemostats, where C:P_R_ and growth rate can be maintained independently, would make the appearance of these multimodal distributions less likely within single-population experiments. Yet, even in chemostats, pure-cultures present heterogeneous growth rates and activities indicating this presumption may be true in some but not all cases^61, 62^. Regardless, there appears to be a stoichiometric trait optimum for each isolate under each condition where the majority of individuals within the population exist (i.e. a single dominant mode). This is expected, as biomass stoichiometry is reflective of the macromolecular composition of the cell, and the relative composition of macromolecules is thought to strongly coupled to growth phase^26, 33^.

### Analysis of natural microbial communities

While we saw small changes in the biomass optima among each lake ecosystem, the trait distributions of both N:P and C:P were markedly different from the stoichiometric trait distribution for the isolate cultures. This was true primarily for the stoichiometric traits containing P. Lake community biomass N:P and C:P had multimodal distributions whereas C:N was consistently unimodal (Figure 3). Average community biomass C:N for all three lakes combined ranged between 4.74 – 4.97; on average lower than what we observed in our isolate experiments and lower than those previously reported (7.3:1) for freshwater bacterial biomass^56^. Due to the conserved^36^ and primarily unimodal distribution of biomass C:N observed here, we suggest a single value for microbial biomass C:N may accurately represent the composition of bacterial biomass in aquatic ecosystem models. However, for the stoichiometric traits of biomass N:P and C:P, multiple modes were present for all three lakes indicating that using a single value to represent these important microbial traits may fail to accurately capture microbial biomass composition in aquatic ecosystem models. Interestingly, the average trait value and trait richness of biomass N:P and C:P of these natural communities was lower than values reported for the isolate experiments (Tables 2 and 3 and Figure 4) and lower than mean trait values previously reported for freshwater bacterial biomass (12:1 and 102:1, respectively)^56^. In effect, the natural communities that were sampled presented a stoichiometric richness that was more similar to the 100:1 C:P_R_ isolate cultures than the 1000:1 C:P_R_ isolate cultures (Figure 4). As lake ecosystems are predicted to be P-limited, this was surprising, especially given the oligotrophic nature of the alpine lake community. Yet, cultured bacterial isolates have been shown previously to exhibit higher biomass C:P and N:P values than average values for freshwater communities^56^ and one possible explanation is that these laboratory experiments occur in the absence of common natural phenomena and ecological interactions such as loss due to sinking, predation, and competition that may serve to increase biomass P.

**Figure 4:**
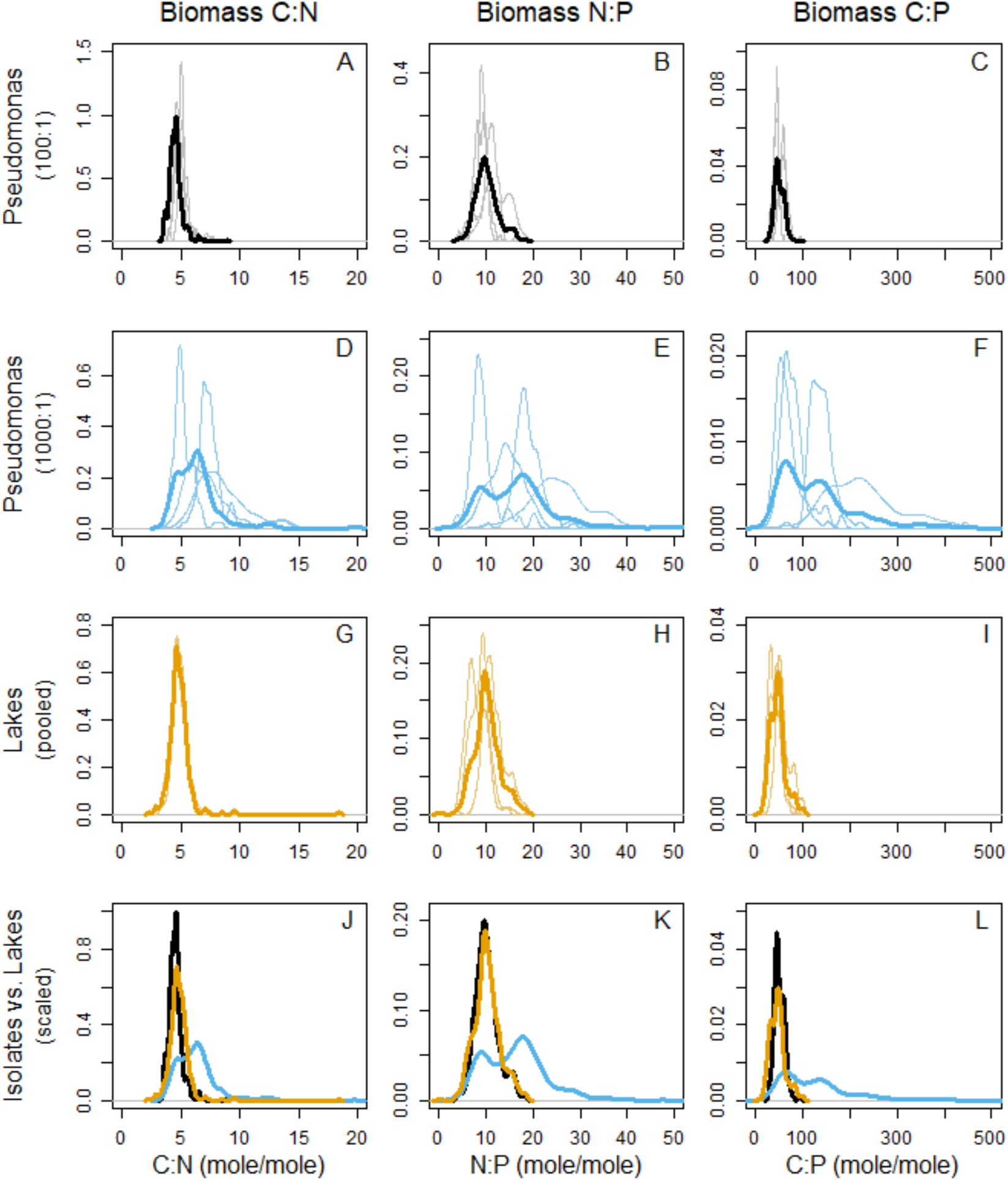
Comparisons of stoichiometric trait distributions between one bacterial population *(Pseudomonas* UND-L18) grown on high-P (panels A-C) and low-P (panels D-F) media to the lake community distributions (G-I). Independent growth phases or lake communities are represented by the lighter, finer lines whereas the aggregated trait distribution is bolder and darker. Aggregated stoichiometric trait distributions are compiled into panels J-L where we present the stoichiometric distributions across the three experiments where black and blue lines indicate the aggregated trait distribution of the high-P and low-P grown *Pseudomonas* cultures, respectively, and the orange line is the aggregated trait distribution from the three lake communities.

The presence of different modes within a stoichiometric trait distribution likely represents ecologically-important microbial life history strategies which may or may not be phylogenetically conserved. As higher peaks in the trait distributions indicate increased abundance of individuals that present that trait value, each of the modes must represent a substantial proportion of the constituents of a population or community. The relative prevalence in multi-mode stoichiometric trait distributions in lake communities compared to the population experiments may reflect a sampling artifact where different microenvironments were sampled and pooled during a sampling event. However, given each sample was collected in a single grab from the surface water of a lake this seems unlikely. Rather, we suggest that each mode in the stoichiometric trait distribution of these natural communities is more likely to be composed of microorganisms with similar ecological strategies. Our results suggest that, within freshwater microbial communities, groups of organisms exist that are adapted or evolved to have a finite number of values for the common stoichiometric traits, which results in multi-modal stoichiometric trait distributions. In additional, that natural bacterial communities that we sampled exhibited a less rich stoichiometric trait distribution compared to cells grown in culture. This suggests that, in the presence of interspecific ecological interactions, microbial cells are likely to only exhibit a realized phenotype and not the full potential of phenotypic plasticity that they may be capable of. As has been suggested previously^64, 65^, models that accurately represent the distribution of trait values within microbial communities are more likely to accurately predict the functional response of microbial communities to changing environmental conditions or across important environmental gradients.

### A new framework in microbial community biomass stoichiometry

The underlying assumption of a trait-based approach is that the traits, rather phylogenetic relationships, are better able to characterize how organisms interact with an ecosystem. Yet, due primarily to technological limitations, trait-based approaches in microbial ecology have a limited resolution in that they have focused on population-or community-level trait signals. We argue that trait-based approaches in microbial ecology could benefit from representations of individual trait values and the use of distributions, rather than averages, in ecosystem simulations. Studying the stoichiometric trait distributions of bacterial populations, under controlled growth conditions, and natural communities, under unknown conditions, creates an opportunity to understand how the environment acts on that distribution, how this distribution influences the community-level trait, and how these community-level traits influence the ecosystem. If communities are evenly distributed around a mean value, as shown in the majority of the isolate growth experiments (Figure 2) and for bacterial biomass C:N, it may be reasonable to use a single estimate for microbial biomass stoichiometry in ecosystem models. On the other hand, if the mean value for a community biomass stoichiometry is really a composite of two or more modes, a single value for a stoichiometric trait may inhibit both understanding and predicting the role of microbial biomass stoichiometry in ecosystems. Regardless, the use of a mean value for microbial traits ignores the phenotypic noise seen in even the most controlled laboratory studies. An individual trait based approach, such as the one presented here, could lead to breakthroughs in our ability to relate community structure to function, forge a link between physiology and genotype, and be used to improve our understanding and modeling of how microorganisms impact principal biogeochemical cycles in a range of ecosystems.

## Experimental Procedures

### Generation and validation of calibration factors for EDS analysis

A full description of the process by which we devised the EDS calibration factors including the EDS analyses used in this study can be found in the Supplemental Materials. In brief, we measured the number of counts in the carbon peak of latex polystyrene beads, corrected for both background signal and area analyzed, to provide a calibration factor to relate C counts to femtomoles of C. We then used this value to generate calibration factors for N and P using microdroplets of three biologically-relevant standards. These calibration factors were found to vary by microdroplet and standard (Supplemental Figure 2) and thus an average calibration factor was utilized for N and P (Supplemental Figure 3 and Supplemental Table 3) throughout.

### Culture and sample preparation of microbial cells

To evaluate the stoichiometric trait distribution of bacterial populations we selected three isolates (a *Pseudomonas* (UND-L18), *Flavobacterium* (N288), and a *Brevundimonas* isolate (N1445)), used in previous studies on microbial stoichiometry^39^, and maintained them using aseptic technique. All isolates were grown on Basal Microbiological Medium (BMM)^39, 6^ with standard media components, with the exception of the trace metal solution where 2.5 ml of this stock solution was added per liter of media. We added carbon as D-glucose to a final concentration in the media of 23.9 mmoles C per liter. We added nitrogen as ammonium chloride to a final concentration of 18.69 mmoles N per liter. We added phosphorus as KH_2_PO_4_ to a final concentration of 239 μM P to provide a C:P_R_ of 100:1 in the high-P media, or to a final concentration of 23.9 μM P for the 1000:1 (low-P media) treatment^39^. We maintained each of the three strains on the high-P media to facilitate rapid growth. Prior to conducting experiments in the low-P media, we transferred each culture repeatedly (>2 times) in the low-P media prior to sampling to ensure we added little to no excess P carry over to the experimental tubes.

Prior to EDS analysis, we harvested cells from the pure-culture experiments by centrifugation (room temperature (RT), 1,950 × *g*, 10 min) and washed 1x with sterile MilliQ water to remove media salts and prevent their deposition onto the grid surface. Following the wash step, we resuspended these samples in 10-200 μl of MilliQ water (dependent on cell density) with a 0.01 % solution of latex beads (1-μm diameter, 64030-15, Polysciences) as an internal standard when desired. We then deposited each cell preparation (10 μl of resuspended cells) directly onto the TEM grid and allowed to dry at room temperature.

### Comparison of bulk biomass C:N:P and EDS biomass C:N:P

To assess whether traditional bulk measurements of biomass C:N:P would provide comparable C:N:P ratios to those obtained from EDS we inoculated (1 % v/v) triplicate 250-ml flasks containing 100:1 C:P_R_ BMM from mid-log cultures grown in identical media. By assessing growth over time and ensuring cultures were sampled during the middle of exponential phase, we attempted to limit the heterogeneity inherent in sampling during non-exponential growth^31^. We selected the *Pseudomonas, Flavobacterium*, and *Brevundimonas* isolates to study bacteria across a range of bacterial phyla (Gammaproteobacteria, Bacteroidetes, and Alphaproteobacteria, respectively). We assessed growth via optical density (OD) at 600 nm and monitored until mid-log phase, the timing of which was determined based on previous growth experiments. At this point, we harvested the cells by centrifugation (RT, 2,456 × *g*, 30 min) and prepared the cells for EDS or bulk analyses.

For EDS analysis, we generated 101 spectra from *Pseudomonas* UND-L18, 92 from *Flavobacterium* N288, and 149 from *Brevundimonas* N1445 and processed the spectra using calibration factors derived from analysis of three biologically-relevant standards (Supplemental Materials). The decision to analyze > 90 cells for each population was based on prior experiments with both low- and high-richness cultures where even the high-richness cultures approached the global mean prior to the measurement of 90 cells (Supplemental Figure 4).

For the bulk analyses, we filtered 100-ml aliquots of cell-containing media from each replicate onto pre-combusted (500 °C for 4 h) GF/F Whatman filters (1825-025, GE Healthcare). After filtration of cultured cells, we rinsed each filter with 10 ml of sterile MilliQ water to remove residual C, N, and P from the media. One filter for each replicate was analyzed for C and N content with a Costech elemental combustion system coupled to a Thermo Scientific Delta V Advantage Isotope Ratio Mass Spectrometer while we used a second filter to determine total phosphorus via an acid persulfate digestion assay^35^ which we measured spectrophotometrically at 880 nm using the molybdenum blue method^35^.

### EDS-assessed stoichiometry across a bacterial growth curve

To demonstrate the ability of EDS to assess interspecific stoichiometric trait distributions in pure cultures, we inoculated the same three isolates analyzed above, in triplicate, into four sets of high- and low-P media to an OD (600 nm) of *ca.* 0.020. We assessed increases in OD hourly or bihourly until the OD plateaued or slowed dramatically, indicating stationary phase. In addition, we collected media hourly or bihourly for the determination of soluble P remaining in the media, as determined by the molybdenum blue method^35^ without prior digestion. We sacrificed cultures at four points across the growth curve (representing lag, early-log, mid-log, and stationary phase) to obtain samples for EDS as growth progressed. These samples thus provided us with snapshots of the stoichiometric trait distribution for each population over time which we mapped to both OD and the availability of P within the media. Sample preparation was performed as previously mentioned, while collection volumes were altered based on the OD of the culture to provide sufficient cell numbers for analysis. When analyzed by EDS, we generated *ca.* 100 spectra for each isolate. Following spectral processing, we normalized the number of spectra for each isolate, across both high- and low-P media, resulting in 90 high-quality spectra being used for *Pseudomonas* UND-L18 and *Brevundimonas* N1445 while 88 spectra were used for *Flavobacterium* N288.

### EDS-assessed stoichiometry from uncultured microbial populations

To assess the distribution of stoichiometric phenotypes in natural mixed bacterioplankton communities, we sampled and analyzed uncultured microbial samples from surface waters of Loch Vale (sampled 09/19/2017, *ca.* 40°17’29.5”N, 105°39’29.8”W), Warren Lake (sampled 10/05/2017, *ca.* 40°31’50.3”N, 105°03’48.2”W), and Sheldon Pond (sampled 12/13/2017, *ca.* 40°34’57.8”N, 105°06’20.7”W) via a two-step filtration process. We removed any large cells and/or particulates via passage through a Whatman GF/C filter (1822-047, Whatman) prior to the concentration of microbial cells via centrifugation (200 ml in 50-ml aliquots, RT, 2,456 × *g*, 30 min) for EDS. For EDS analysis, we generated 218 spectra from the Loch Vale sample, 203 from the Warren Lake sample, and 160 spectra from the Sheldon Pond sample with sufficient quality (background- and film-subtracted C counts ≥ 3,000) to enable accurate C:N:P calculations.

### TPDs and calculation of FD components

Trait probability density functions were generated using the density function in the R software platform^67^ with the smoothing bandwidth set to 80 % for each TPD presented within this work. We performed calculations of trait diversity including trait richness, evenness, and divergence by using the TPD package (TPD: Methods for Measuring Functional Diversity Based on Trait Probability Density, Version 1.0.0)^1^ in the R software platform^67^. To compare characteristics of each distribution across the cultured and uncultured microbial communities we analyzed each strain, time point, and lake community simultaneously. We selected an alpha value of 0.95 for each trait to make the analyses less sensitive to outliers. We set the trait range for biomass C:N to 0 − 20, biomass N:P to 0 − 100, and biomass C:P to 0 − 1000 to cover the full range of each trait value seen in our experimental data following trimming via the 0.95 alpha value. In addition, we set the number of divisions within those ranges to provide one division per 0.1 change in molar ratio.

## Acknowledgements

This work is a product of NSF IOS award DEB#1456959 awarded to EKH. The authors would like to thank the Cotner Lab (University of Minnesota) for supplying the freshwater bacterial isolates used in the growth experiments. The Central Instrument Facility, Department of Chemistry (Colorado State University) provide instrument time and support during the method development portion of the current work. The current version of this manuscript was improved by comments provided by Adam Martiny (University of California, Irvine)

